## Supplementary Materials for "Uniform bacterial genetic diversity along the gut"

#### Table of Contents

##### Supplementary Tables

Supplementary Table 1. Sequencing reads per sample
Supplementary Table 2. Results of pathways analysis
Supplementary Table 3. $\pi$ estimates
Supplementary Table 4. Variance in strain frequencies within and between cages
Supplementary Table 5. Allele frequencies of SNVs experiencing extreme frequency changes
Supplementary Table 6. Allele frequencies of SNVs experiencing extreme frequency changes in healthy humans

##### Supplementary Figures

Supplementary Figure 1. Differential abundance of bacterial families along the gut
Supplementary Figure 2. Functional pathway differences along the gut.
Supplementary Figure 3. Species considered for each analysis
Supplementary Figure 4. Inoculum $\pi$ and average mouse $\pi$
Supplementary Figure 5. Relative strain frequency along the guts of humanized mice
Supplementary Figure 6. Variance in major strain frequency partitioned between gut region, mouse, and cage in humanized mice
Supplementary Figure 7. Variance in strain frequencies within and between cages
Supplementary Figure 8. Evolutionary changes along the guts of humanized mice
Supplementary Figure 9. Increase in taxonomic diversity and change in community membership along the length of the guts of conventional mice.
Supplementary Figure 10. Relative strain frequency along the guts of conventional mice
Supplementary Figure 11. Variance in major strain frequency partitioned between gut region, mouse, and cage in conventional mice.
Supplementary Figure 12. Relative strain frequency along the guts of healthy humans
Supplementary Figure 13. Evolutionary changes along the guts of healthy humans.
Supplementary Figure 14. Inferring strain frequency of *Bacteroides vulgatus* strains
Supplementary Figure 15. Supervised strain frequency inference of *Bacteroides uniformis* strains

### Supplementary Tables

[Supplementary Table 1 attached as .xlsx file]

**Supplementary Table 1. Sequencing reads per sample in humanized mice and conventional mice studies.** This workbook includes the following sheets:

***Humanized Mice.*** The total number of metagenomic for all humanized mouse samples. Shown at the bottom of the table is the median sequencing depth across all mouse samples.

***Conventional Mice.*** The total number of metagenomic reads for all conventional mouse samples.

[Supplementary Table 2 attached as .xlsx file]

**Supplementary Table 2. Pathways Significantly Associated with Gut Location.** The raw p-values and Benjamini-Hochberg adjusted p-values (q-value) for 86 pathways significantly associated with gut location at $q < 0.05$ measured by coefficient for binary covariate representing either small or large intestine.

[Supplementary Table 3 attached as .xlsx file]

**Supplementary Table 3. $\pi$ estimates.** $\pi$, a measure of nucleotide diversity, was estimated for 30 species in all mouse samples and the inoculum sample for which the species had at least 500,000 loci that had a coverage $\geq 4$.

[Supplementary Table 4 attached as an .xlsx file]

**Supplementary Table 4. Variance in strain frequencies within and between cages.**

For each species with detected co-colonization of two strains, the variance of the inferred dominant strain frequency was computed among groups of samples. This workbook includes the following sheets:

***strain_freq_variance_btw_cage***. Variance in dominant strain frequency observed at the same gut region (duodenum, jejunum, ileum, cecum or colon) in mice of different cages.

***strain_freq_variance_wthn_cage***. Variance in dominant strain frequency observed at the same gut region among mice co-housed in either cage 1, 2, or 3.

[Supplementary Table 5 attached as .xlsx file]

**Supplementary Table 5. Allele frequencies of SNVs experiencing extreme frequency changes.** This workbook includes the following sheets:

***All SNV freq changes.*** Allele frequencies of all SNVs detected as having gone from an allele frequency of $f \leq 0.2$ to $f \geq 0.8$ between any pair of samples. SNVs are annotated according to species, the types of sample pairs they were observed changing between ("Observed in"), contig, position in the reference genome ("site_pos"), PATRIC ID ("gene_id"), gene annotations lifted from the PATRIC database ("gene_description"), and codon degeneracy ("variant_type," where 1D indicates that the site is 1-fold degenerate such that any nucleotide difference will result in an amino acid change, and 4D indicates that the site is 4-fold degenerate such that any nucleotide difference will not result in an amino acid change). Subsequent columns indicate the sample in which the allele frequencies are calculated. An allele frequency value of NA indicates that the nucleotide site did not meet the minimum read coverage requirement of 20 reads or more in that sample.

***All within-host SNV freq changes.*** The 4 within-host SNVs are annotated with the species where the SNV was detected, the subject in which the change was observed to occur between the small and large intestine, average allele frequencies in the small and large intestine, and the magnitude of difference between the small and large intestine.

[Supplementary Table 6 attached as .xlsx file]

**Supplementary Table 6. Allele frequencies of SNVs experiencing extreme frequency changes in healthy humans.** This workbook includes the following sheets:

***Change between regions.*** Metadata for each SNV detected as having gone from an allele frequency of $f \leq 0.2$ to $f \geq 0.8$ between any pair of samples taken at different regions and at the same time point. Shown in each row is the respective time point, Capsules ID corresponding to a specific gut region (Methods) of each sample, allele frequencies of each sample, allele frequency change, and description of the gene in which the SNV was detected. The grey bars in the third column indicate SNVs in close proximity that are likely linked.

***Change between timepoints.*** Metadata for each SNV detected as having gone from an allele frequency of $f \leq 0.2$ to $f \geq 0.8$ between any pair of samples taken at different time points. Shown in each row are the pairs of samples in which the SNV change was detected and their respective time points, Capsules ID corresponding to a specific gut region (Methods), allele frequencies, allele frequency change, and description of the gene in which the SNV was detected. The grey bars in the third column indicate SNVs in close proximity that are likely linked.

***Change between time points summary.*** Summary data for the SNVs depicted in *Change between time points* worksheet. The minimum and maximum allele frequency change between time points are shown for each SNV detected as having extreme changes between time points. The grey bars in the third column indicate SNVs in close proximity that are likely linked.

### Supplementary Figures

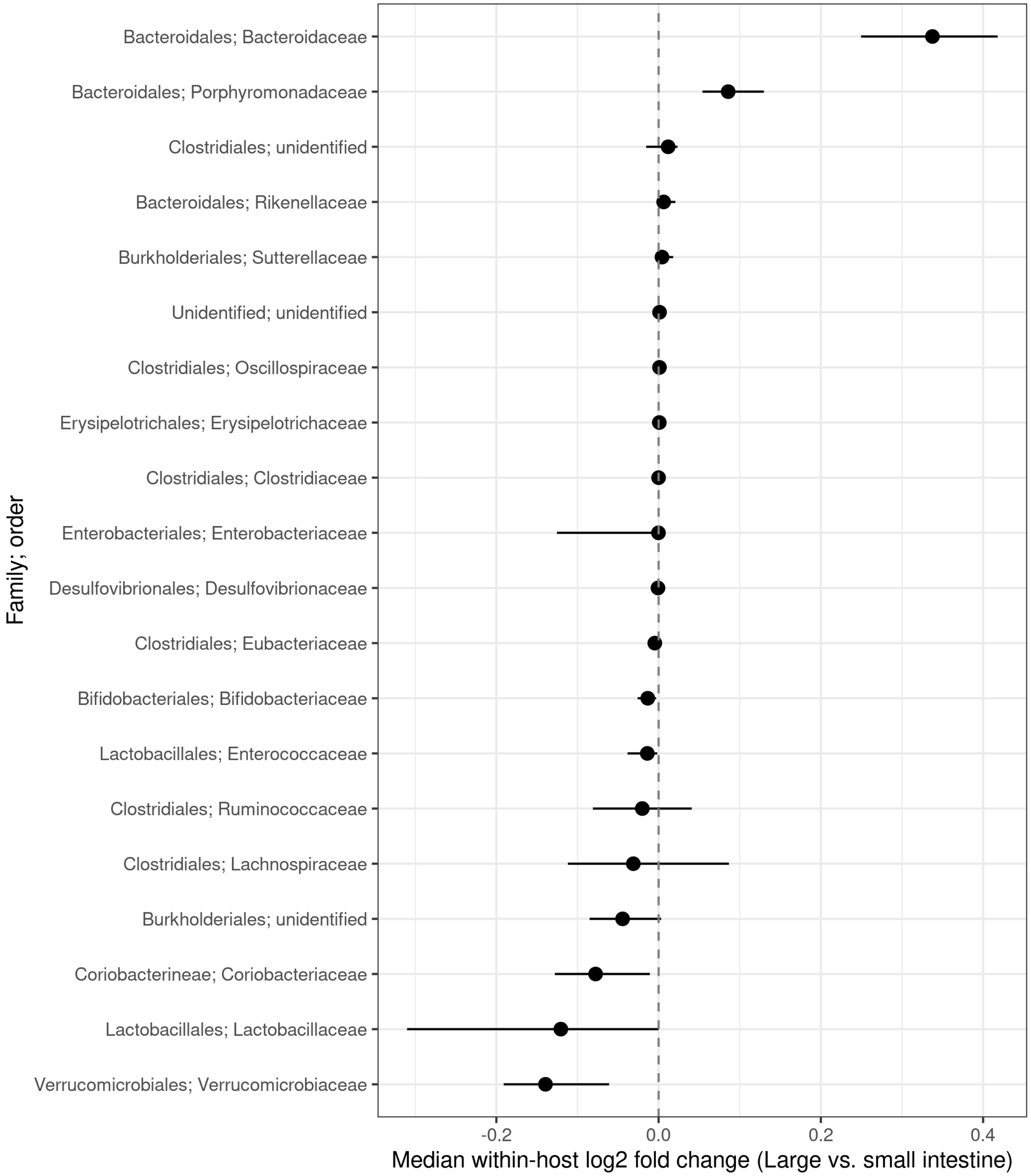


**Supplementary Figure 1. Families have differential relative abundance along the gut.** A paired Wilcoxon signed-rank test was used to estimate median log2 fold change between paired family relative abundance values corresponding to the large and small intestine of the same mice (**Methods**). Taxa labels represent *Order*; *Family*, and are ordered from highest to lowest log2fold change. Positive log2 fold change values indicate families that are enriched in the large intestine, whereas negative log2 fold change values indicate families that are enriched in the small intestine.

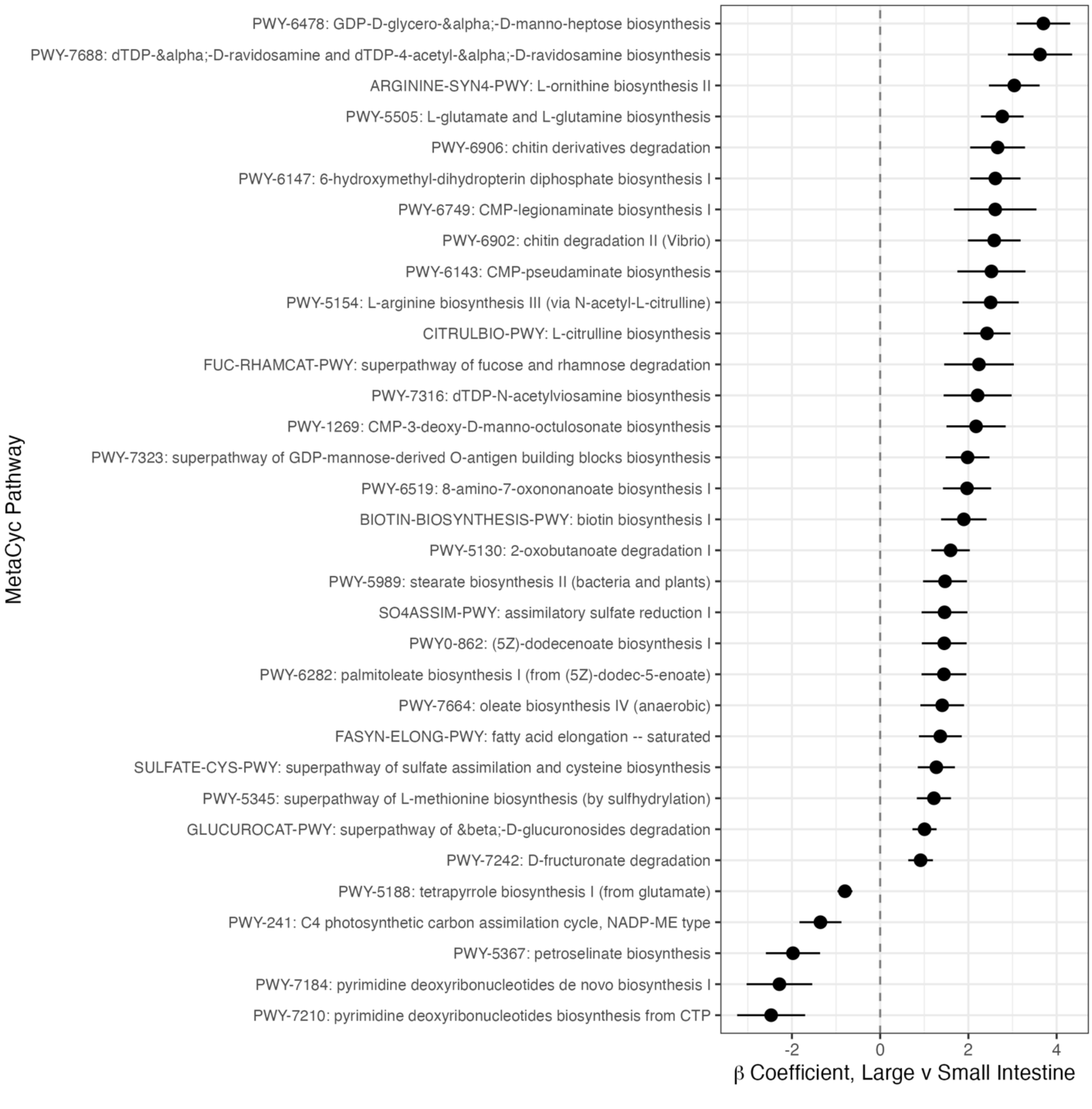


**Supplementary Figure 2. Functional pathway differences along the gut.** Estimated coefficients for the effect of gut location on $\log_2$ relative abundances from a MaAsLIN 3 multivariate linear model. Shown are 34 MetaCyc pathways with highly significant association ($q < 0.001$), none of which were associated with cage or read count ($q > 0.1$). Shown is the 95% confidence interval of the coefficient estimates. Gut location was defined as a binary variable indicating either the large or small intestine. Positive coefficients indicate families that are enriched in the large intestine, whereas negative coefficients indicate families that are enriched in the small intestine. Error bars represent 95% confidence intervals of the coefficient estimates.

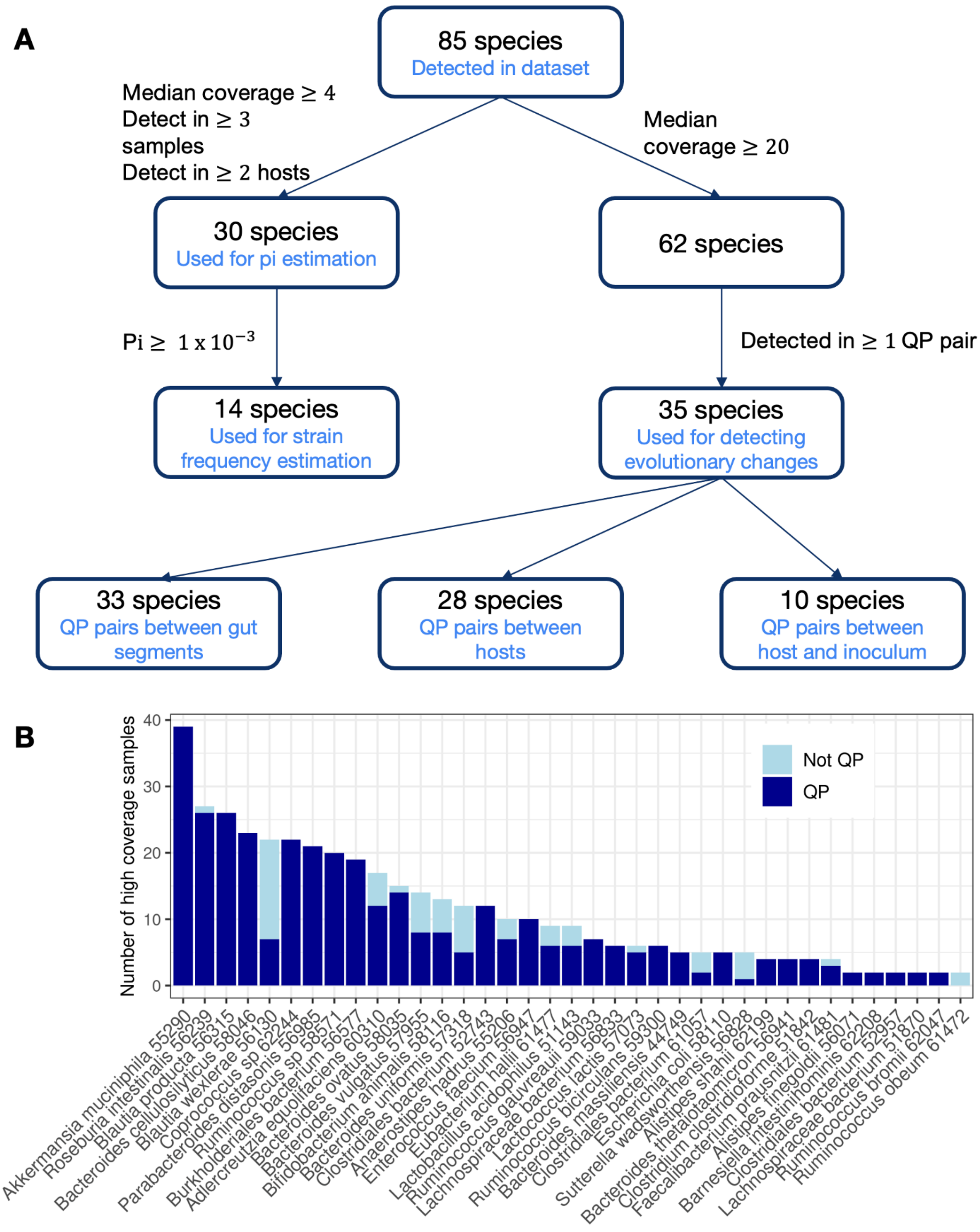

**Supplementary Figure 3. Species considered for each analysis. (A)** Species requirements for assessing within-species diversity with pi, inferring strain frequencies, and assessing evolutionary changes within hosts, between hosts and between inoculum and host. **(B)** Number of quasi-phaseable (QP) and non-quasi-phaseable (non-QP) pairs for all species in the dataset that have at least two high coverage samples (i.e., median coverage ≥ 20).

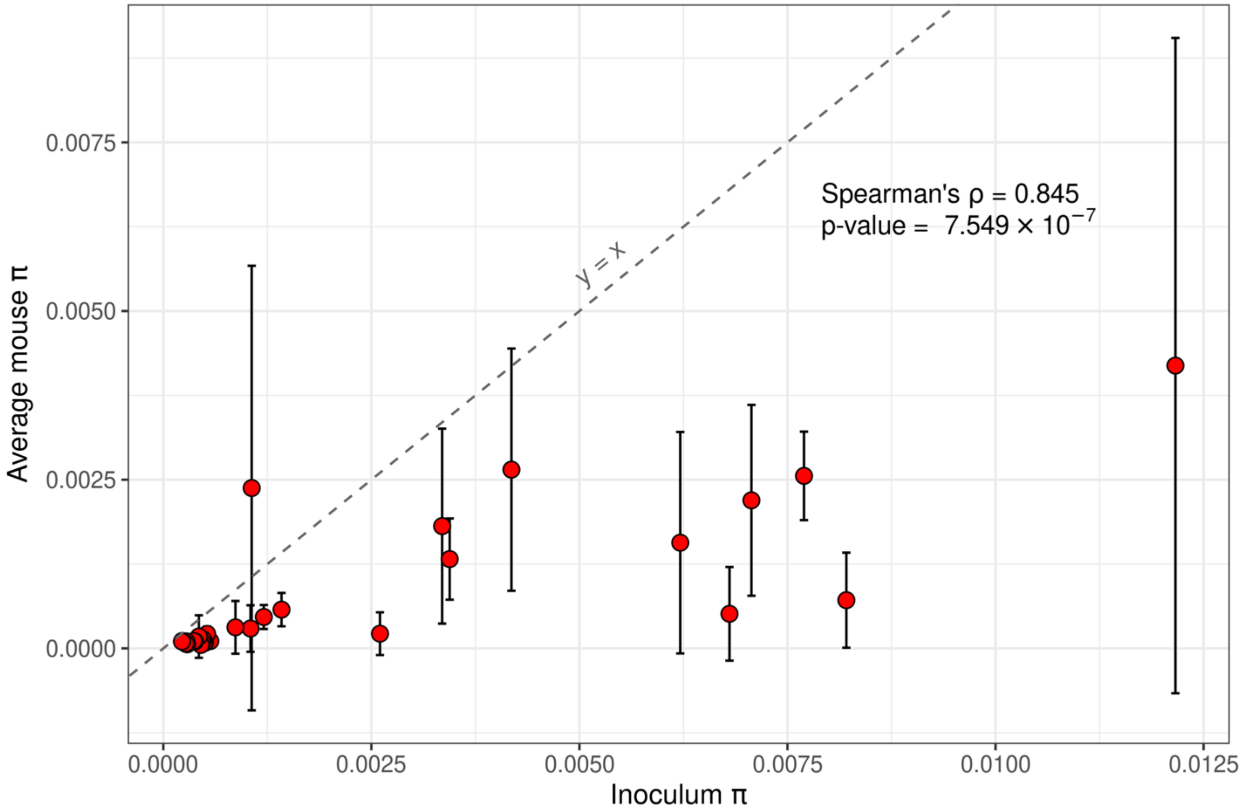


**Supplementary Figure 4. Inoculum $\pi$ and average mouse $\pi$.** Nucleotide diversity ($\pi$) observed in the inoculum (x axis) and average nucleotide diversity observed across mouse samples (y axis) is visualized for the 30 most abundant and prevalent species in the humanized mouse cohort (**Methods**). Error bars represent the standard deviation of $\pi$ values observed in mouse samples. The dashed grey line represents the identity line (y = x).

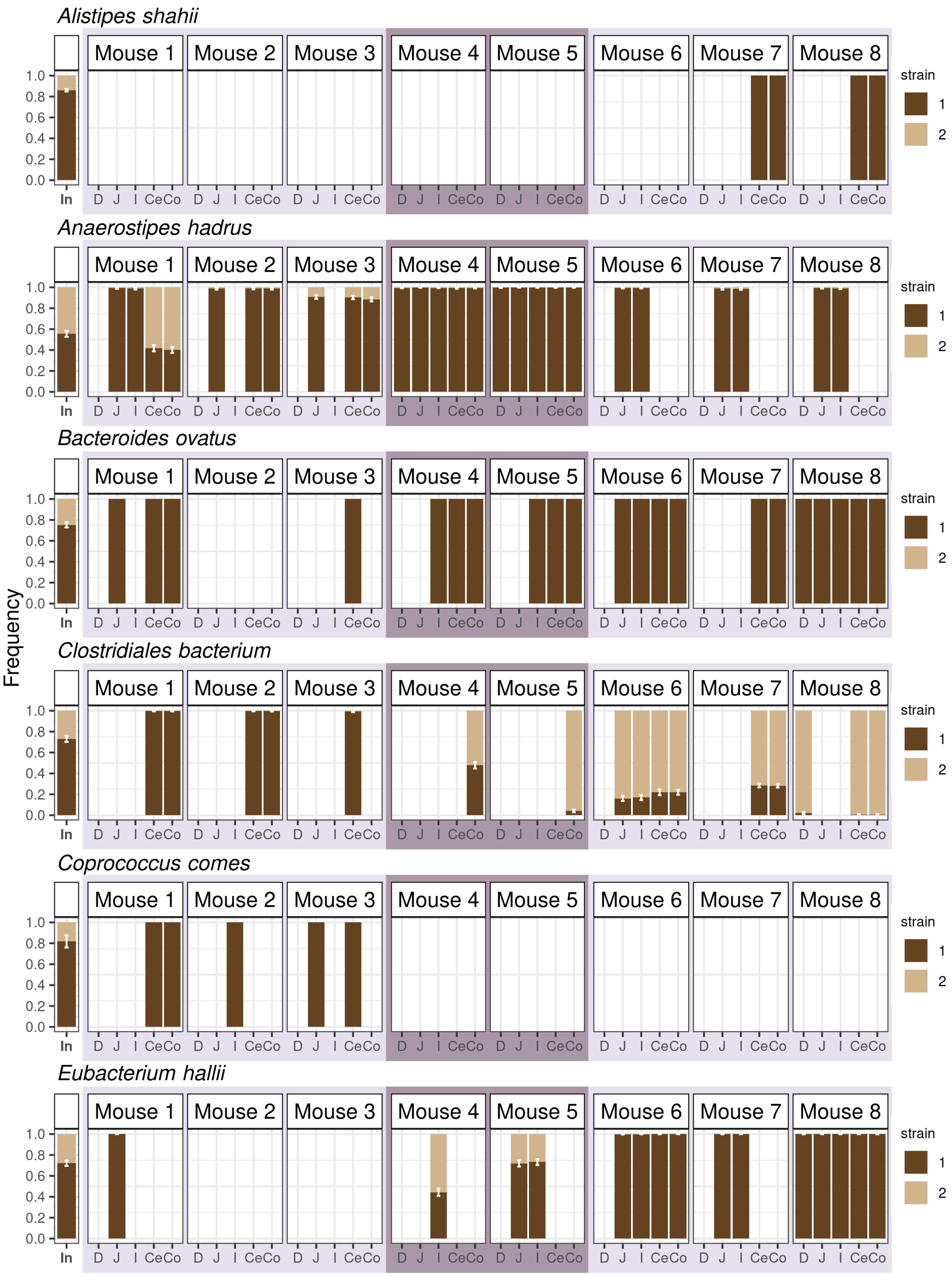

Frequency
Alistipes shahii
Anaerostipes hadrus
Bacteroides ovatus
Clostridiales bacterium
Coprococcus comes
Eubacterium hallii
Mouse 1
Mouse 2
Mouse 3
Mouse 4
Mouse 5
Mouse 6
Mouse 7
Mouse 8
1.0
0.8
0.6
0.4
0.2
0.0
In
D J I CeCo
strain
1
2

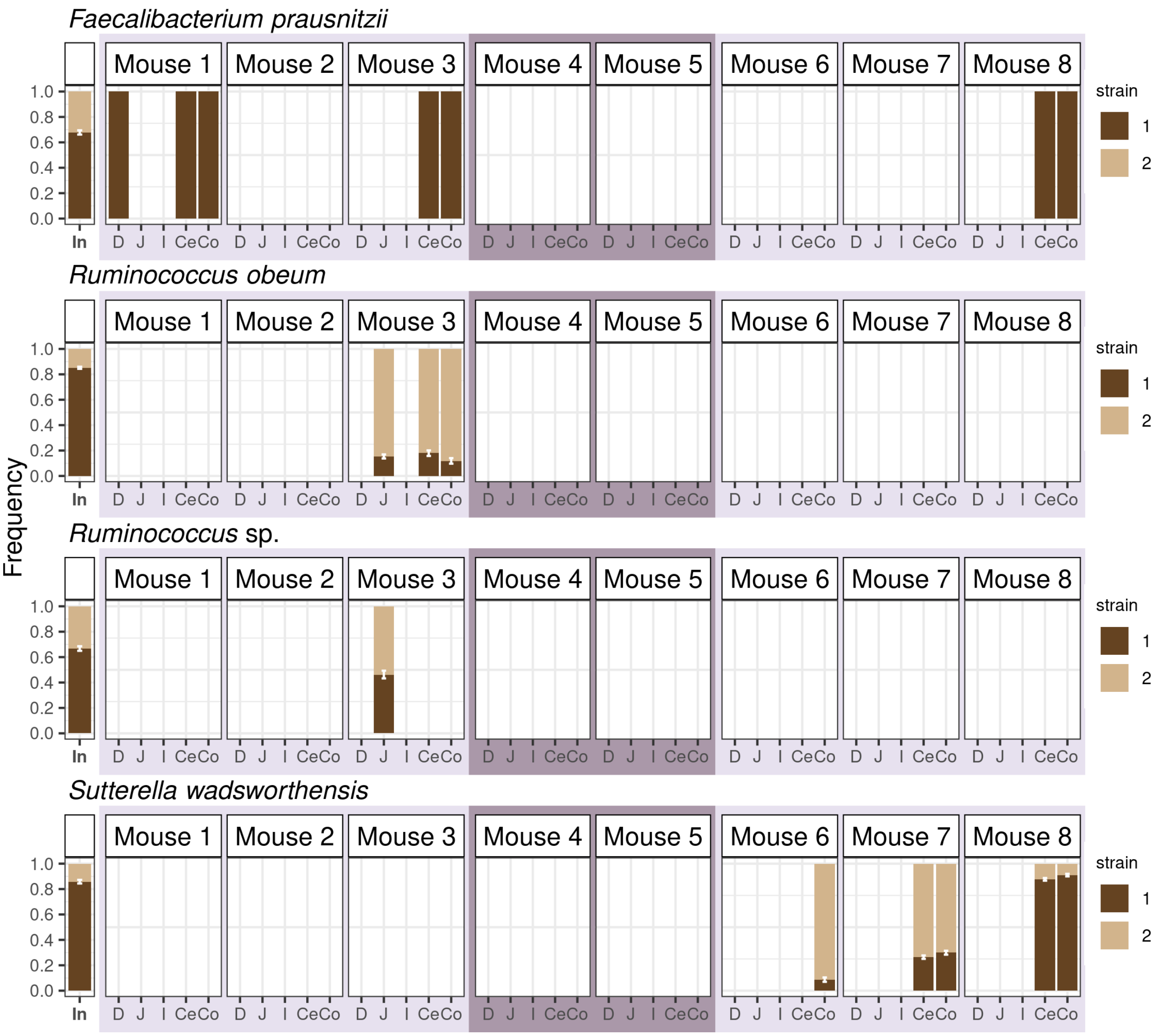


**Supplementary Figure 5. Relative strain frequency along the guts of humanized mice.** Strain frequency of oligo-colonizing strains was inferred across all samples for 10 species that had $\pi \geq 1 \times 10^{-3}$ in the inoculum. Strain frequency is indicated on the y-axis, with error bars representing the 95% confidence intervals for the inferred strain frequency (**Methods**). Cages 1-3 are delineated with alternating light and dark purple boxes. Strain frequencies of the four species not shown here can be found in **Figure 4** of the main text.

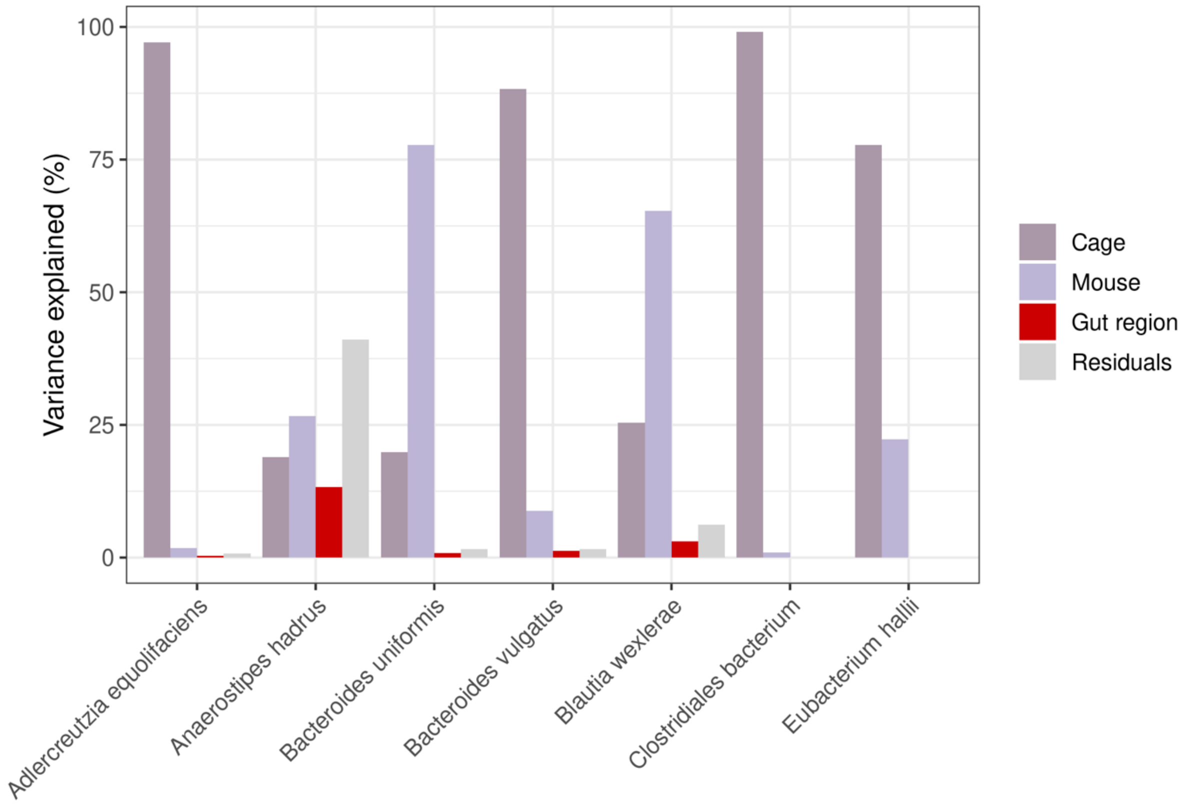


**Supplementary Figure 6. Variance in major strain frequency partitioned between gut region, mouse, and cage in humanized mice.** ANOVA was used to quantify the amount of variance in major strain relative frequency explained by "cage", "mouse", and "gut region" in the seven species for which enough high coverage samples were available to test the effect of all three variables (**Methods**). Residuals of the ANOVA represent unexplained variance.

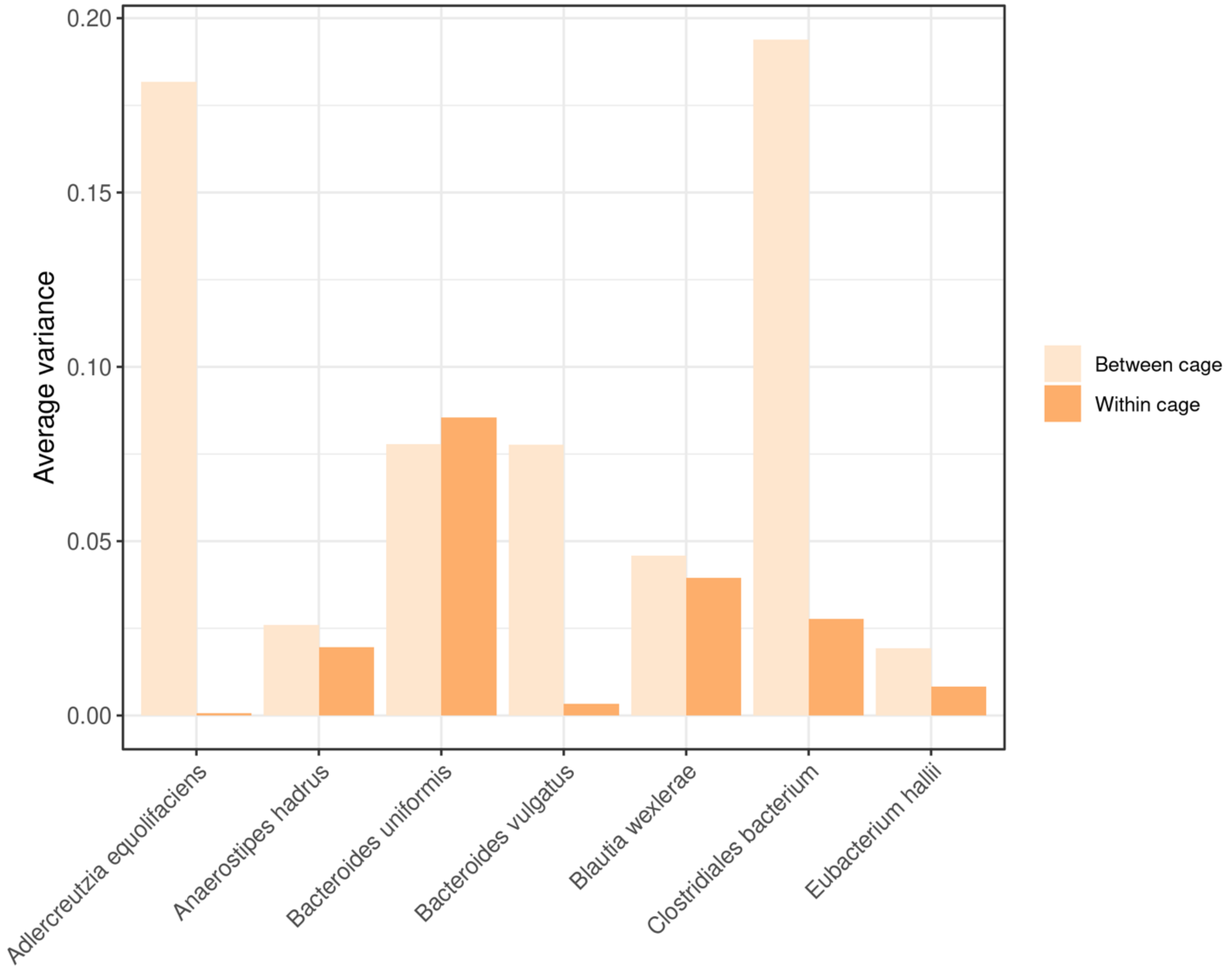


**Supplementary Figure 7. Variance in strain frequencies within and between cages.** Variance in major strain frequency was measured for samples belonging to the same gut region in different mice within the same cage ("Within cage," dark orange) and between different cages ("Between cage," light orange). Within cage bars represent the average within cage variance across the three cages and five gut regions, while between cage bars represent between cage variance averaged across the five gut regions.

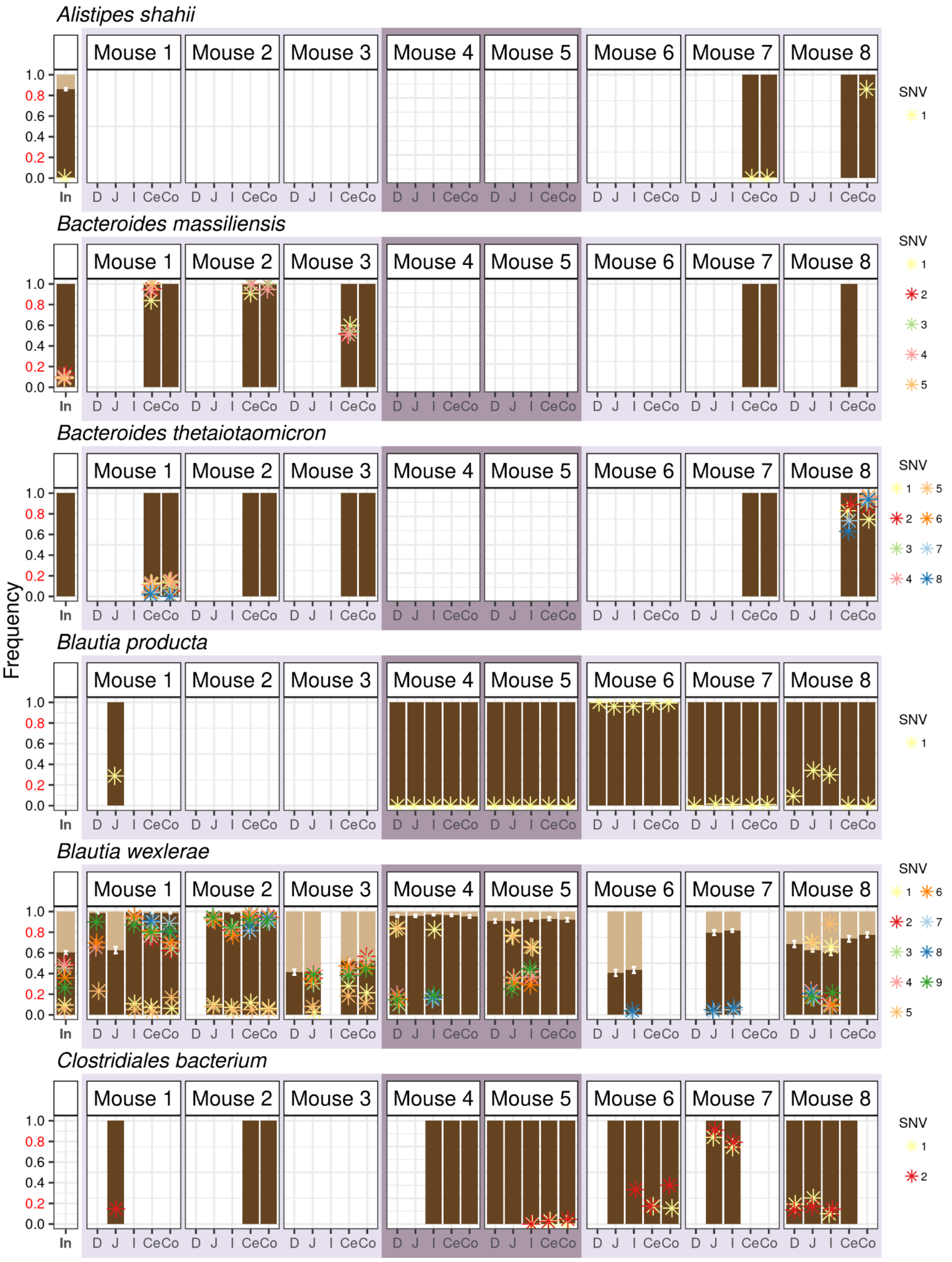
Alistipes shahii
Bacteroides massiliensis
Bacteroides thetaiotaomicron
Blautia producta
Blautia wexlerae
Clostridiales bacterium
Mouse 1
Mouse 2
Mouse 3
Mouse 4
Mouse 5
Mouse 6
Mouse 7
Mouse 8
Frequency
1.0
0.8
0.6
0.4
0.2
0.0
In
D J I CeCo
SNV
1 2 3 4 5 6 7 8 9

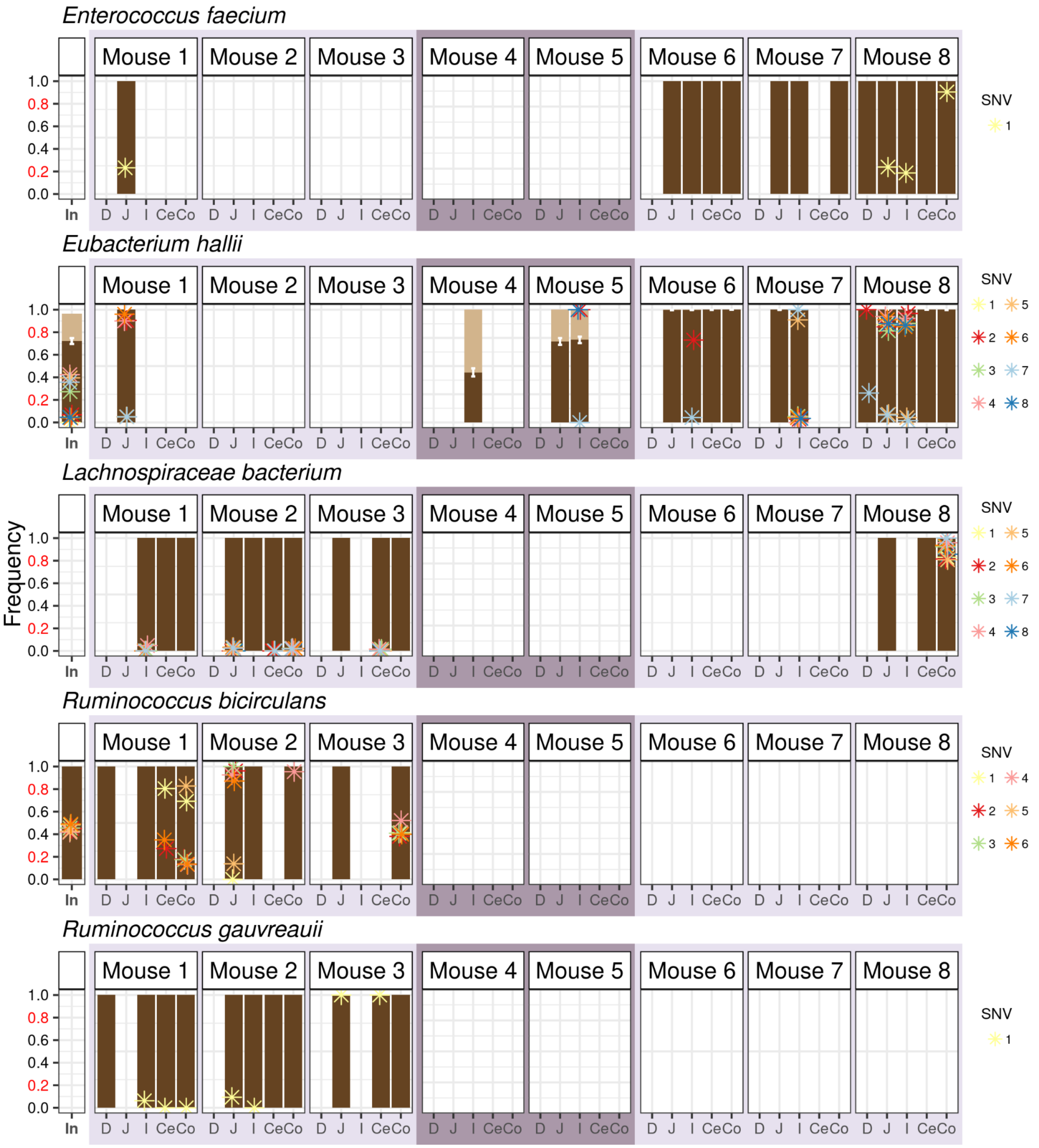


**Supplementary Figure 8. Evolutionary changes along the guts of humanized mice.** SNVs in 13 species underwent extreme allele frequency changes (i.e., $f \leq 0.2$ to $f \geq 0.8$) between pairs of QP samples. Allele frequencies for these SNVs were calculated in all samples for which the loci of interest had a coverage $D \geq 20$. Asterisks represent the allele frequency of a given SNV, with each SNV within a species having its own unique color. Samples lack asterisks for particular SNV when those loci do not have adequate coverage to infer allele frequency. When multiple strains of the same species are present, frequencies of co-colonizing strains are represented as dark and light shades of brown (see **Supplementary Figure 4** for legend).

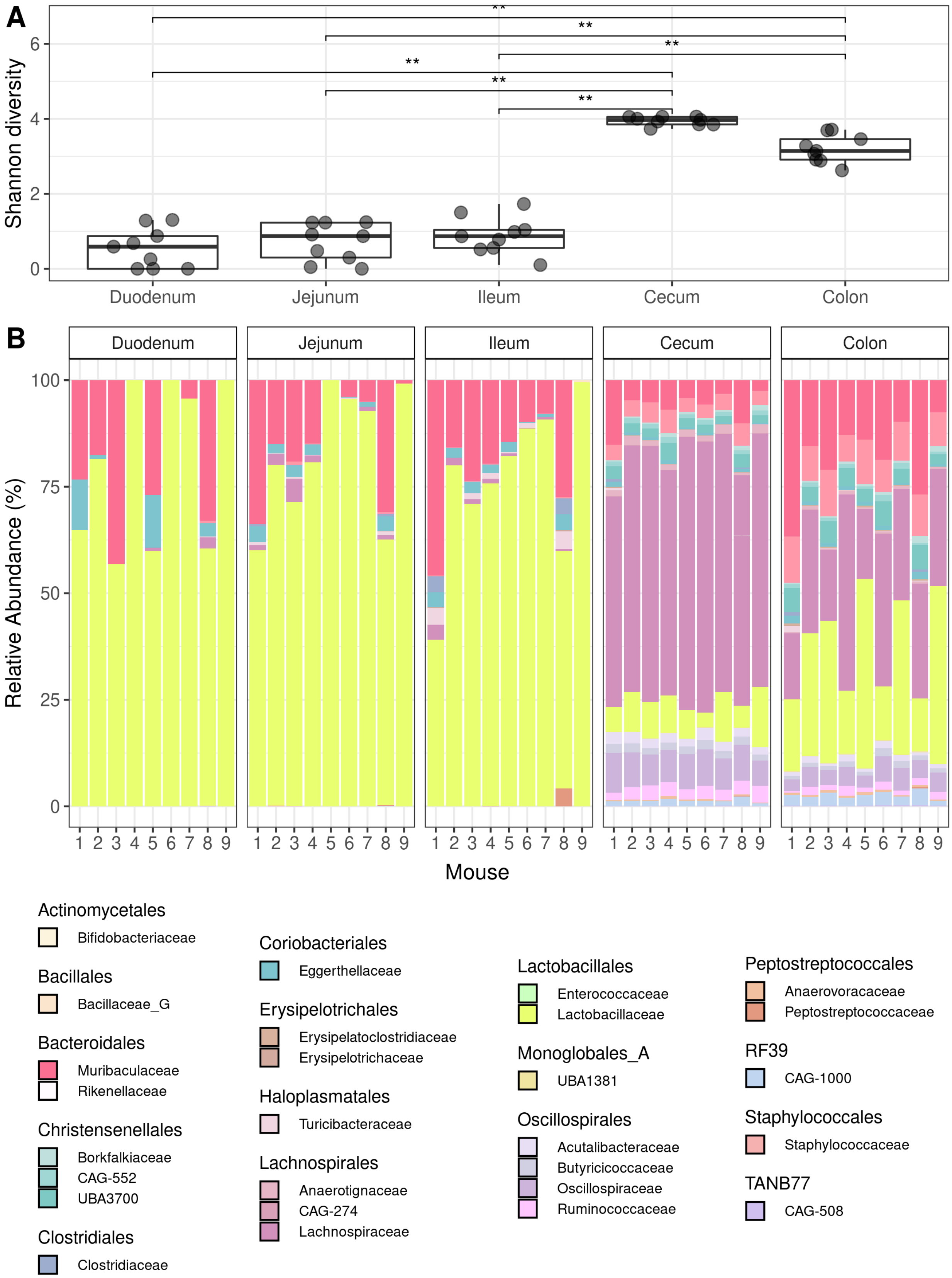
A
B
Shannon diversity
Duodenum
Jejunum
Ileum
Cecum
Colon
Relative Abundance (%)
Mouse
Actinomycetales
Bifidobacteriaceae
Bacillales
Bacillaceae_G
Bacteroidales
Muribaculaceae
Rikenellaceae
Christensenellales
Borkfalkiaceae
CAG-552
UBA3700
Clostridiales
Clostridiaceae
Coriobacteriales
Eggerthellaceae
Erysipelotrichales
Erysipelatoclostridiaceae
Erysipelotrichaceae
Haloplasmatales
Turicibacteraceae
Lachnospirales
Anaerotignaceae
CAG-274
Lachnospiraceae
Lactobacillales
Enterococcaceae
Lactobacillaceae
Monoglobales_A
UBA1381
Oscillospirales
Acutalibacteraceae
Butyricicoccaceae
Oscillospiraceae
Ruminococcaceae
Peptostreptococcales
Anaerovoracaceae
Peptostreptococcaceae
RF39
CAG-1000
Staphylococcales
Staphylococcaceae
TANB77
CAG-508

**Supplementary Figure 9. Increase in taxonomic diversity and change in community membership along the length of the guts of conventional mice. (A)** Alpha diversity (Shannon Index) estimates for bacteria in different regions of the gut. Two-sided Wilcoxon rank sum tests were performed between all possible pairs of small intestinal versus large intestinal samples (** indicates $p \leq 0.01$). **(B)** Relative abundance of bacterial families in the five gut regions and inoculum. Bacterial families are grouped by order in the legend.

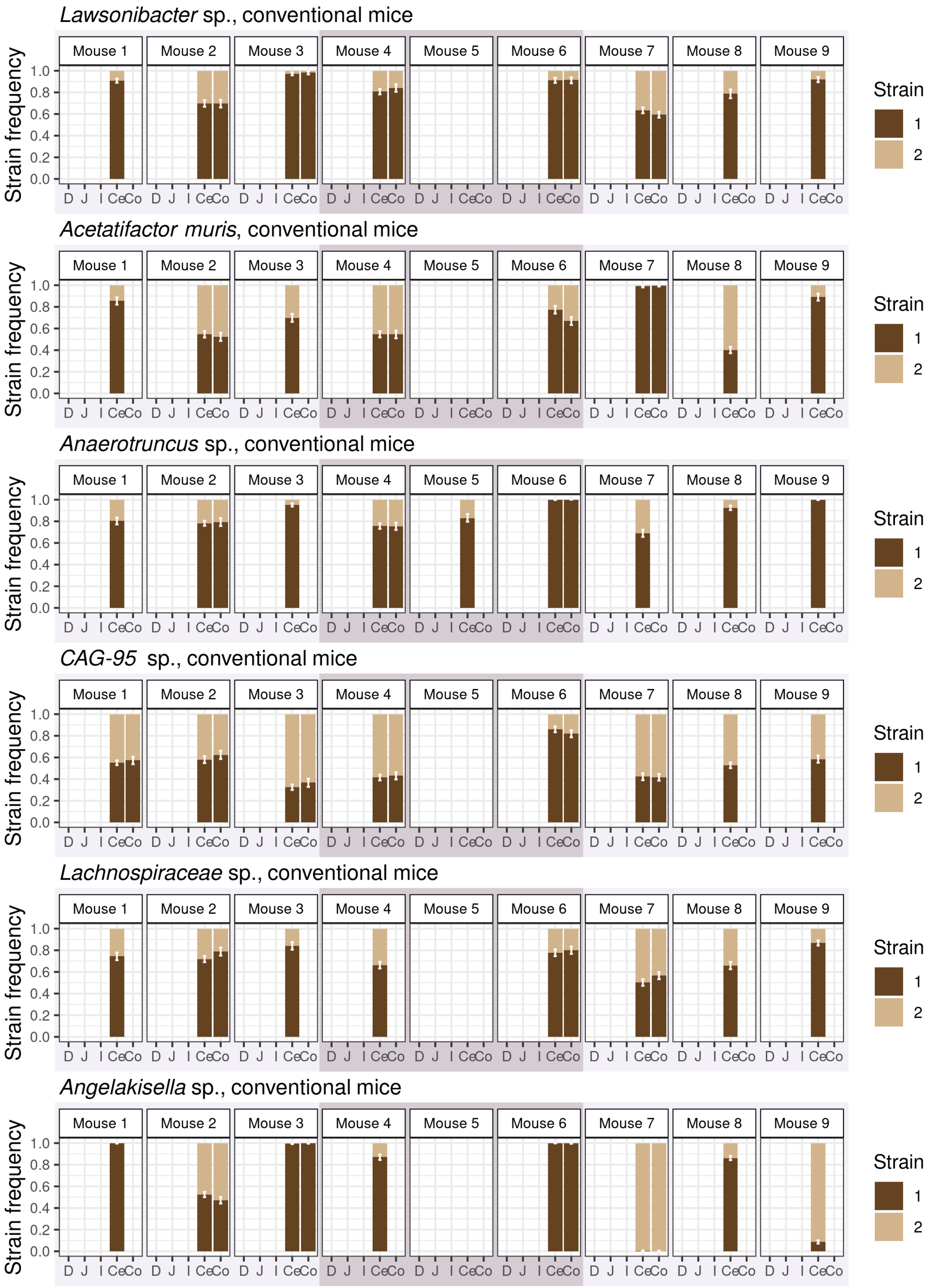
Lawsonibacter sp., conventional mice
Acetatifactor muris, conventional mice
Anaerotruncus sp., conventional mice
CAG-95 sp., conventional mice
Lachnospiraceae sp., conventional mice
Angelakisella sp., conventional mice
Mouse 1
Mouse 2
Mouse 3
Mouse 4
Mouse 5
Mouse 6
Mouse 7
Mouse 8
Mouse 9
Strain frequency
0.0
0.2
0.4
0.6
0.8
1.0
D J I CeCo
Strain
1
2

**Supplementary Figure 10. Relative strain frequency along the guts of conventional mice.** Strain frequency of oligo-colonizing strains was inferred across all conventional mouse samples for six species shown here. Strain frequency is indicated on the y-axis, with error bars representing the 95% confidence intervals for the inferred strain frequency (**Methods**). Cages 1-3 are delineated with alternating light and dark purple boxes. Strain frequencies of the one species with multiple colonizing strains not shown here can be found in **Figure 6A** of the main text.

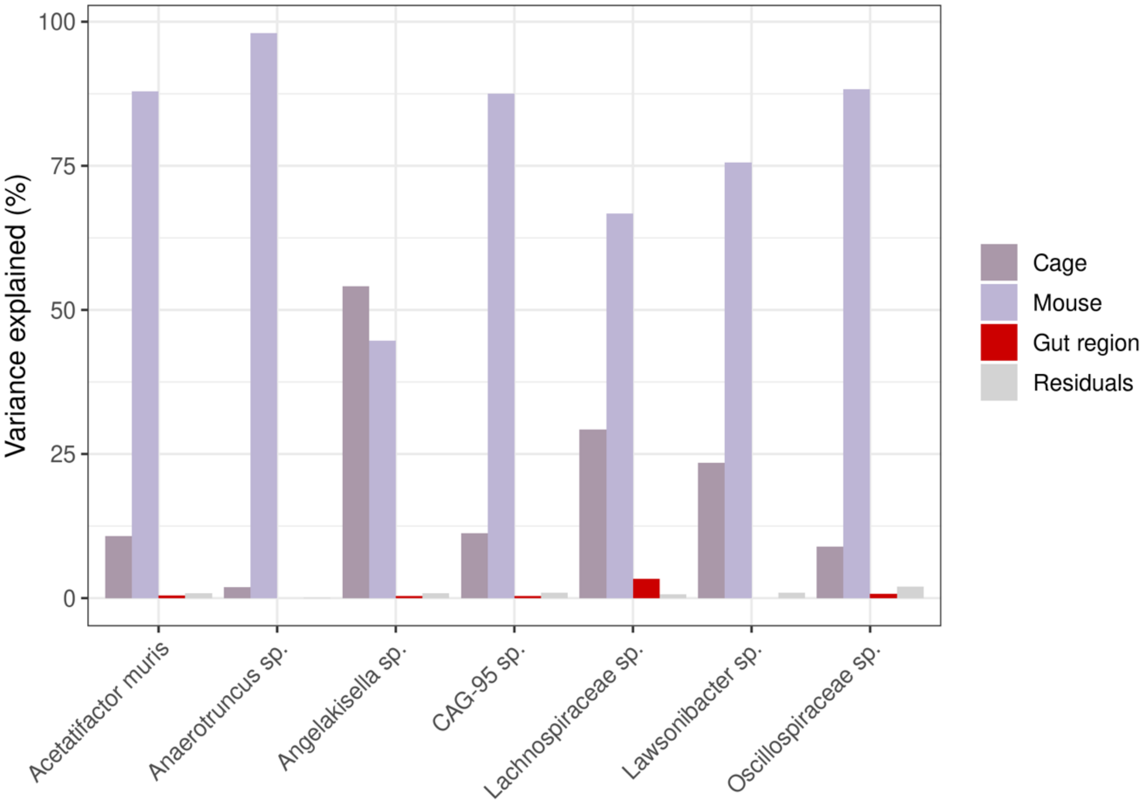


**Supplementary Figure 11. Variance in major strain frequency partitioned between gut region, mouse, and cage in conventional mice.** ANOVA was used to quantify the amount of variance in major strain relative frequency explained by “cage”, “mouse”, and “gut region” in the seven species for which enough high coverage samples were available to test the effect of all three variables (**Methods**). Residuals of the ANOVA represent unexplained variance.

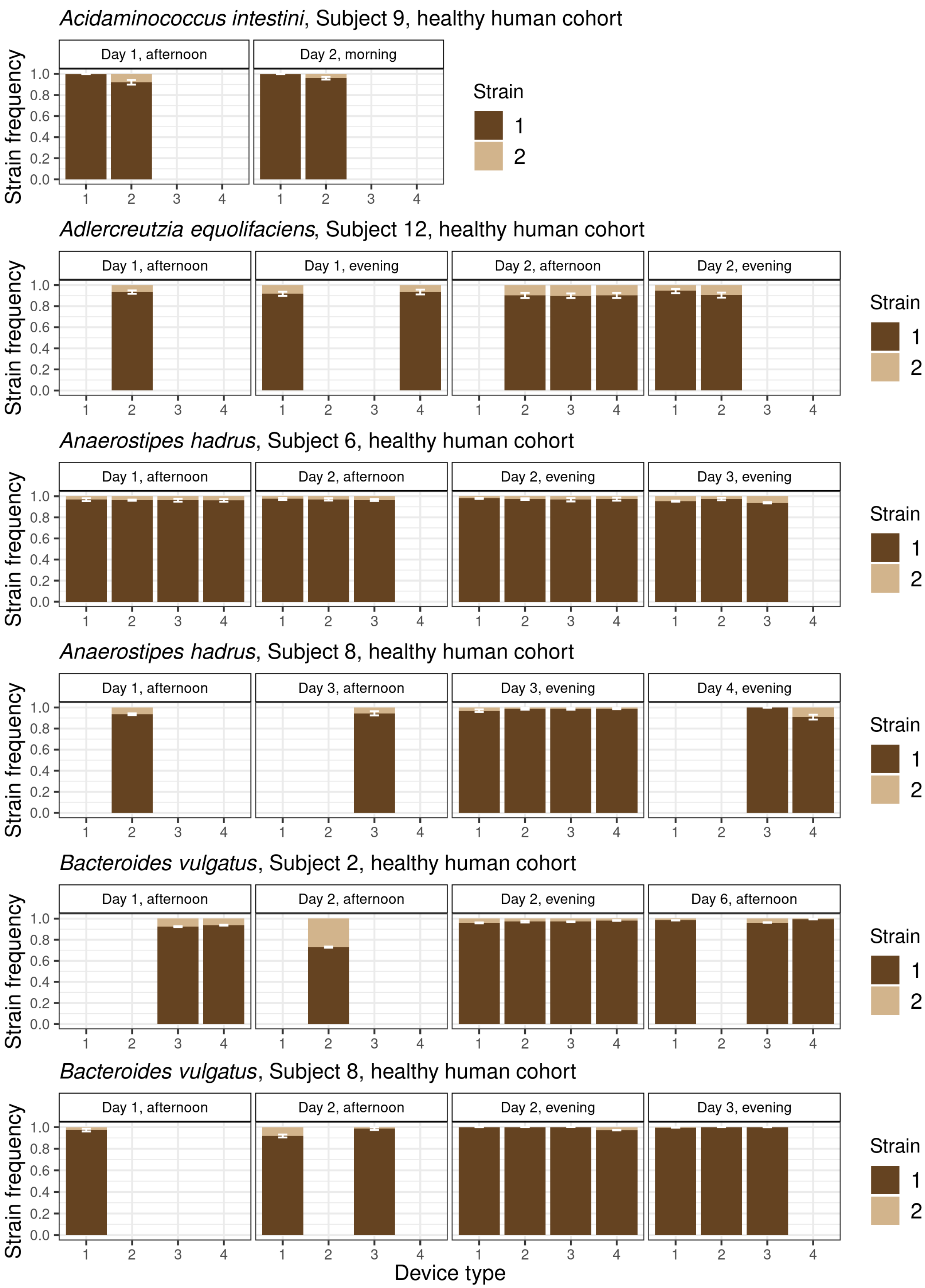
Acidaminococcus intestini, Subject 9, healthy human cohort
Day 1, afternoon
Day 2, morning
Adlercreutzia equolifaciens, Subject 12, healthy human cohort
Day 1, afternoon
Day 1, evening
Day 2, afternoon
Day 2, evening
Anaerostipes hadrus, Subject 6, healthy human cohort
Day 1, afternoon
Day 2, afternoon
Day 2, evening
Day 3, evening
Anaerostipes hadrus, Subject 8, healthy human cohort
Day 1, afternoon
Day 3, afternoon
Day 3, evening
Day 4, evening
Bacteroides vulgatus, Subject 2, healthy human cohort
Day 1, afternoon
Day 2, afternoon
Day 2, evening
Day 6, afternoon
Bacteroides vulgatus, Subject 8, healthy human cohort
Day 1, afternoon
Day 2, afternoon
Day 2, evening
Day 3, evening
Strain frequency
1.0
0.8
0.6
0.4
0.2
0.0
1
2
3
4
Strain
1
2
Device type

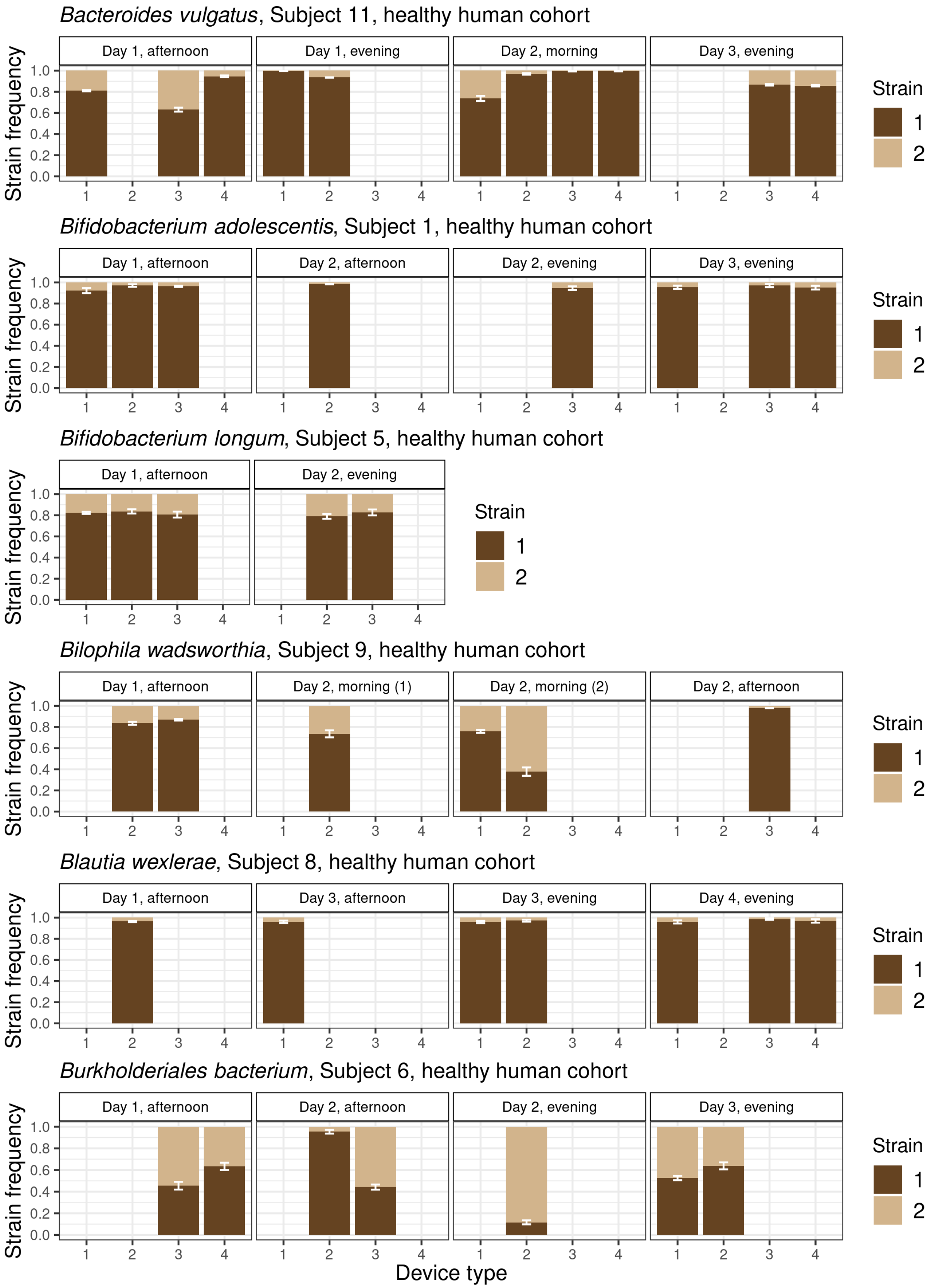

*Bacteroides vulgatus*, Subject 11, healthy human cohort
Day 1, afternoon
Day 1, evening
Day 2, morning
Day 3, evening
*Bifidobacterium adolescentis*, Subject 1, healthy human cohort
Day 1, afternoon
Day 2, afternoon
Day 2, evening
Day 3, evening
*Bifidobacterium longum*, Subject 5, healthy human cohort
Day 1, afternoon
Day 2, evening
*Bilophila wadsworthia*, Subject 9, healthy human cohort
Day 1, afternoon
Day 2, morning (1)
Day 2, morning (2)
Day 2, afternoon
*Blautia wexlerae*, Subject 8, healthy human cohort
Day 1, afternoon
Day 3, afternoon
Day 3, evening
Day 4, evening
*Burkholderiales bacterium*, Subject 6, healthy human cohort
Day 1, afternoon
Day 2, afternoon
Day 2, evening
Day 3, evening
Strain frequency
0.0
0.2
0.4
0.6
0.8
1.0
1
2
3
4
Device type
Strain
1
2

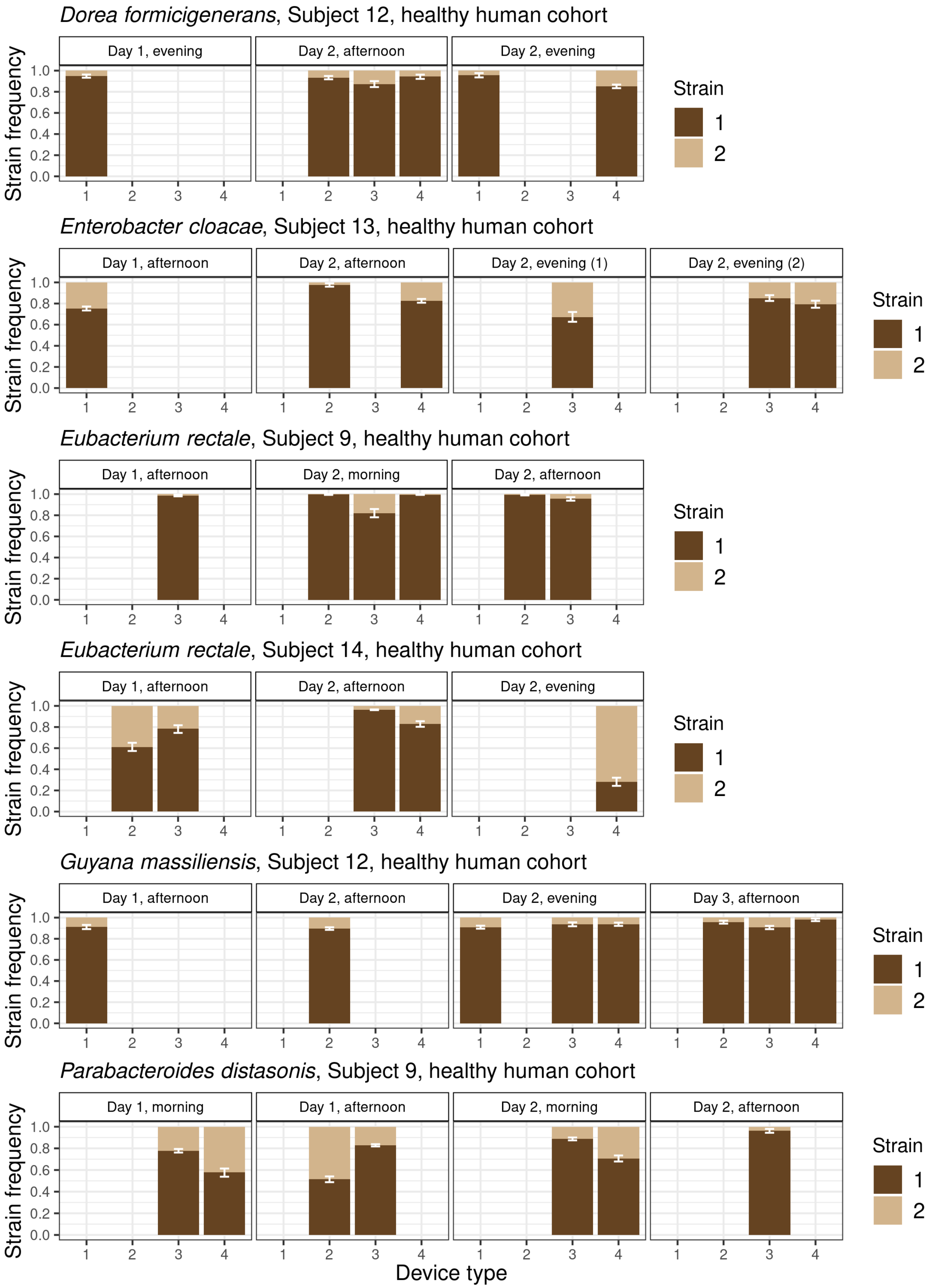
Dorea formicigenerans, Subject 12, healthy human cohort
Day 1, evening
Day 2, afternoon
Day 2, evening
Enterobacter cloacae, Subject 13, healthy human cohort
Day 1, afternoon
Day 2, afternoon
Day 2, evening (1)
Day 2, evening (2)
Eubacterium rectale, Subject 9, healthy human cohort
Day 1, afternoon
Day 2, morning
Day 2, afternoon
Eubacterium rectale, Subject 14, healthy human cohort
Day 1, afternoon
Day 2, afternoon
Day 2, evening
Guyana massiliensis, Subject 12, healthy human cohort
Day 1, afternoon
Day 2, afternoon
Day 2, evening
Day 3, afternoon
Parabacteroides distasonis, Subject 9, healthy human cohort
Day 1, morning
Day 1, afternoon
Day 2, morning
Day 2, afternoon
Strain frequency
0.0
0.2
0.4
0.6
0.8
1.0
1
2
3
4
Device type
Strain
1
2

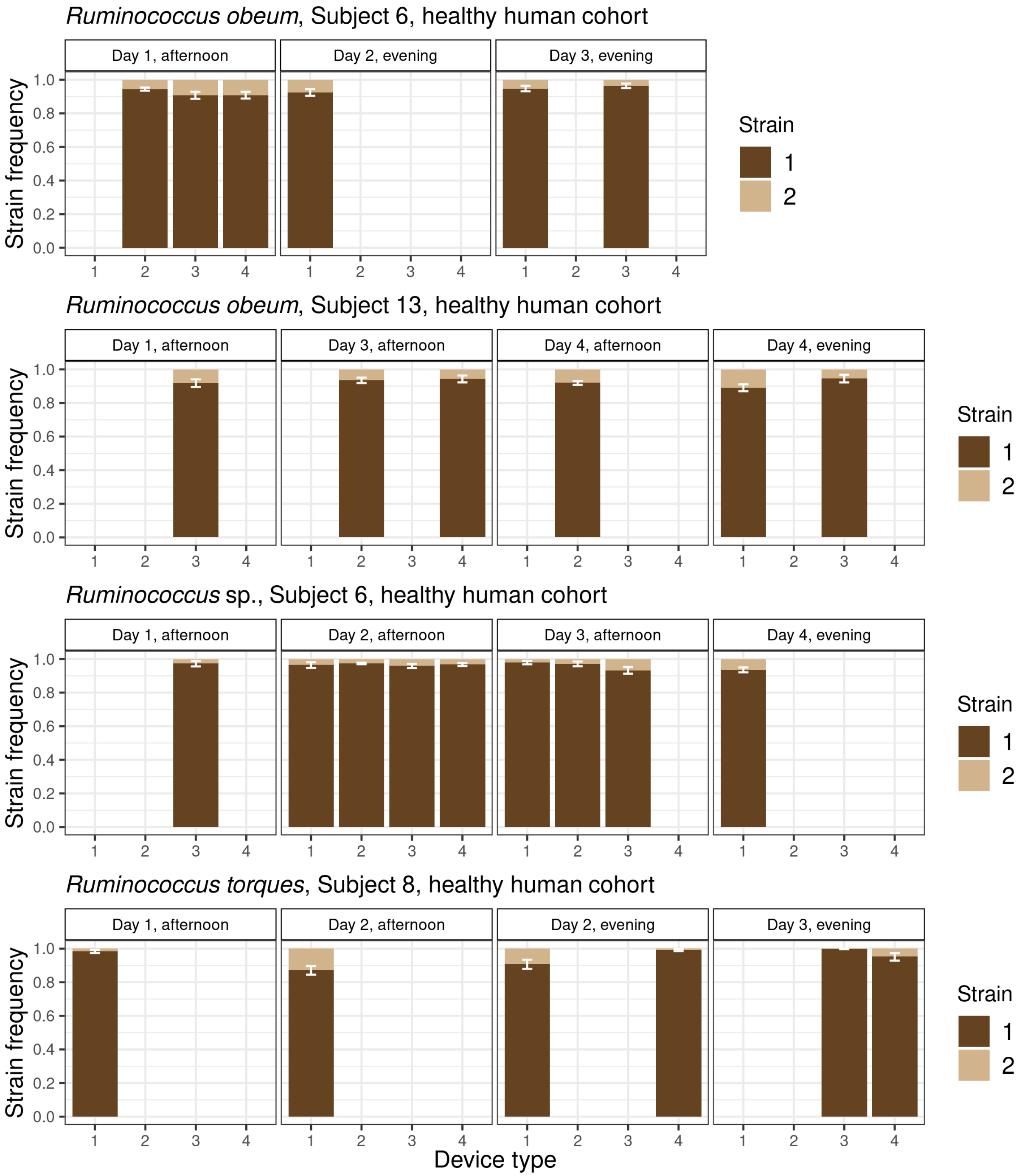


**Supplementary Figure 12. Relative strain frequency along the guts of healthy humans.** Strain frequency of species with oligo-colonizing strains was inferred in a cohort of healthy human subjects. These subject swallowed capsule devices that collected luminal contents along their guts, where device type 1 targeted the pyloric sphincter to the upper small intestine, device type 2 targeted the upper to mid-small intestine, device type 3 targeted the mid- to lower small intestine, and device type 4 targeted the lower small intestine into the ascending colon. Visualized here are the 23 species *x* host pairs (representing 18 species across 11 subjects) which met prevalence filters (**Methods**). If strain frequency was inferred in more than four timepoints, four were manually chosen for visualization here. Strain frequency is indicated on the y-axis, with error bars representing the 95% confidence intervals for the inferred strain frequency (**Methods**). Timepoints represent the time at which capsules were swallowed, with days being relative to the first timepoint plotted and times of day being coarsened into “morning” (before 12 pm PST), “afternoon” (after 12pm and before 8 pm PST), and “evening” (after 8 pm PST) bins. Strain frequencies of the one species in a subject with multiple colonizing strains not shown here can be found in **Figure 6D** of the main text.

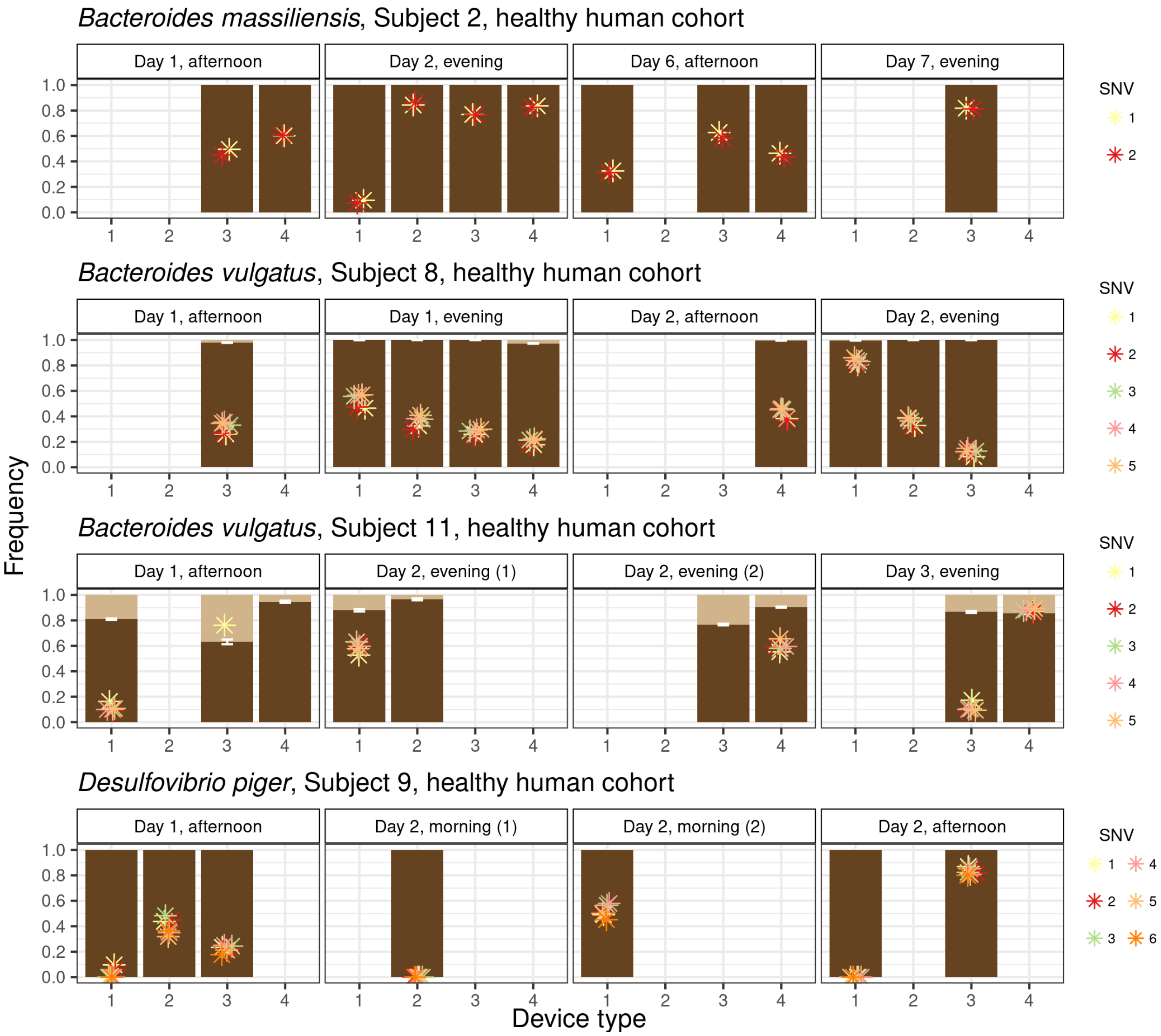


**Supplementary Figure 13. Evolutionary changes along the guts of healthy humans.** We visualized SNVs undergoing extreme allele frequency changes (i.e., $f \leq 0.2$ to $f \geq 0.8$) in at least timepoint in species x host that had sufficient temporal and spatial data (At least two timepoints with at least two device types per timepoint). These filters yielded 18 SNVs detected 4 species x host pairs, representing 3 unique species across 4 unique hosts. Allele frequencies for these SNVs were calculated in all samples for which the loci of interest had a coverage $D \geq 20$. Asterisks represent the allele frequency of a given SNV, with each SNV within a species having its own unique color. Samples lack asterisks for SNVs when those SNVs do not have adequate coverage to infer allele frequency. When multiple strains of the same species are present, frequencies of co-colonizing strains are represented as dark and light shades of brown (See **Supplementary Figure 14** for legend). Timepoints represent the time at which capsules were swallowed, with days being relative to the first timepoint plotted and times of day being coarsened into "morning" (before 12 pm PST), "afternoon" (after 12pm and before 8 pm PST), and "evening" (after 8 pm PST) bins.

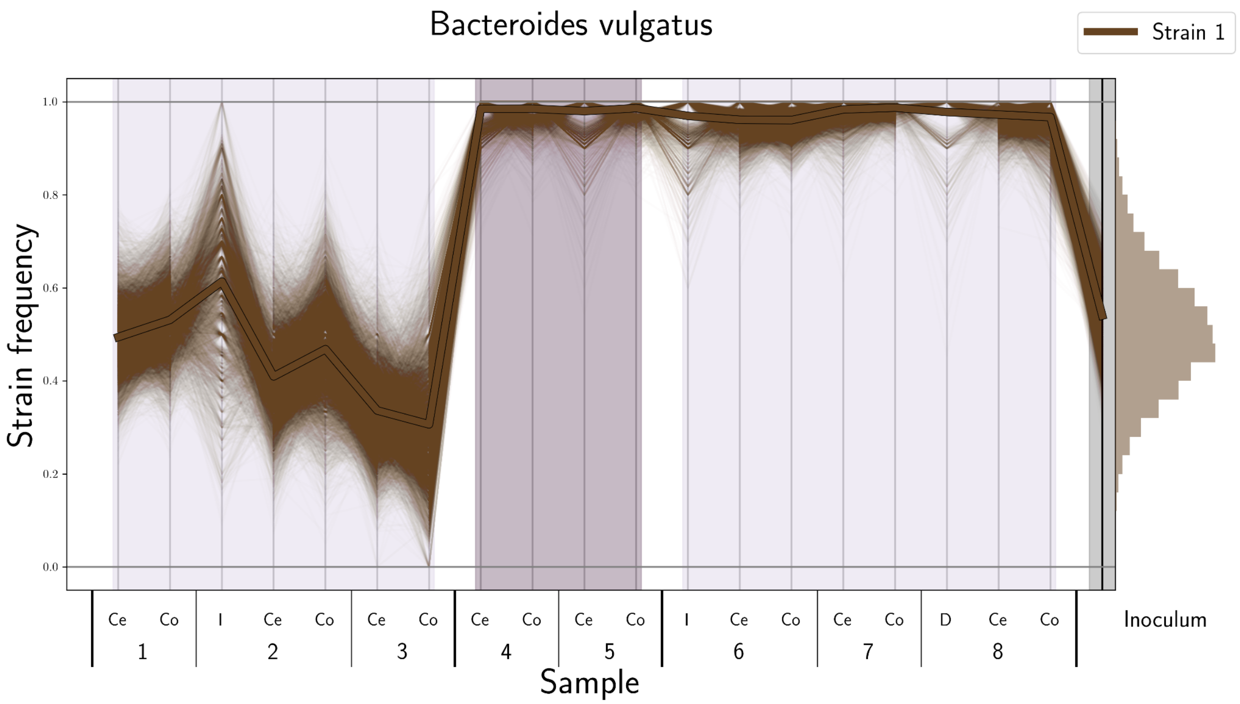


**Supplementary Figure 14. Inferring strain frequency of *Bacteroides vulgatus* strains.** Strain frequencies were inferred for *B. vulgatus* (and other species) by clustering loci into large groups of SNVs that display highly correlated allele frequencies (thin brown lines) across samples (indicated by labels on the x axis; "D" indicates duodenum, "I" indicates ileum, "Ce" indicates cecum, and "Co" indicates colon; numbers indicate mouse identity). The strain frequency was inferred to be the mean of the inferred clusters (thick brown line). In the case of *B. vulgatus*, one cluster was inferred, distinguishing two strains. The histogram to the right of the plot represents the distribution of allele frequencies of clustered SNVs in the inoculum.

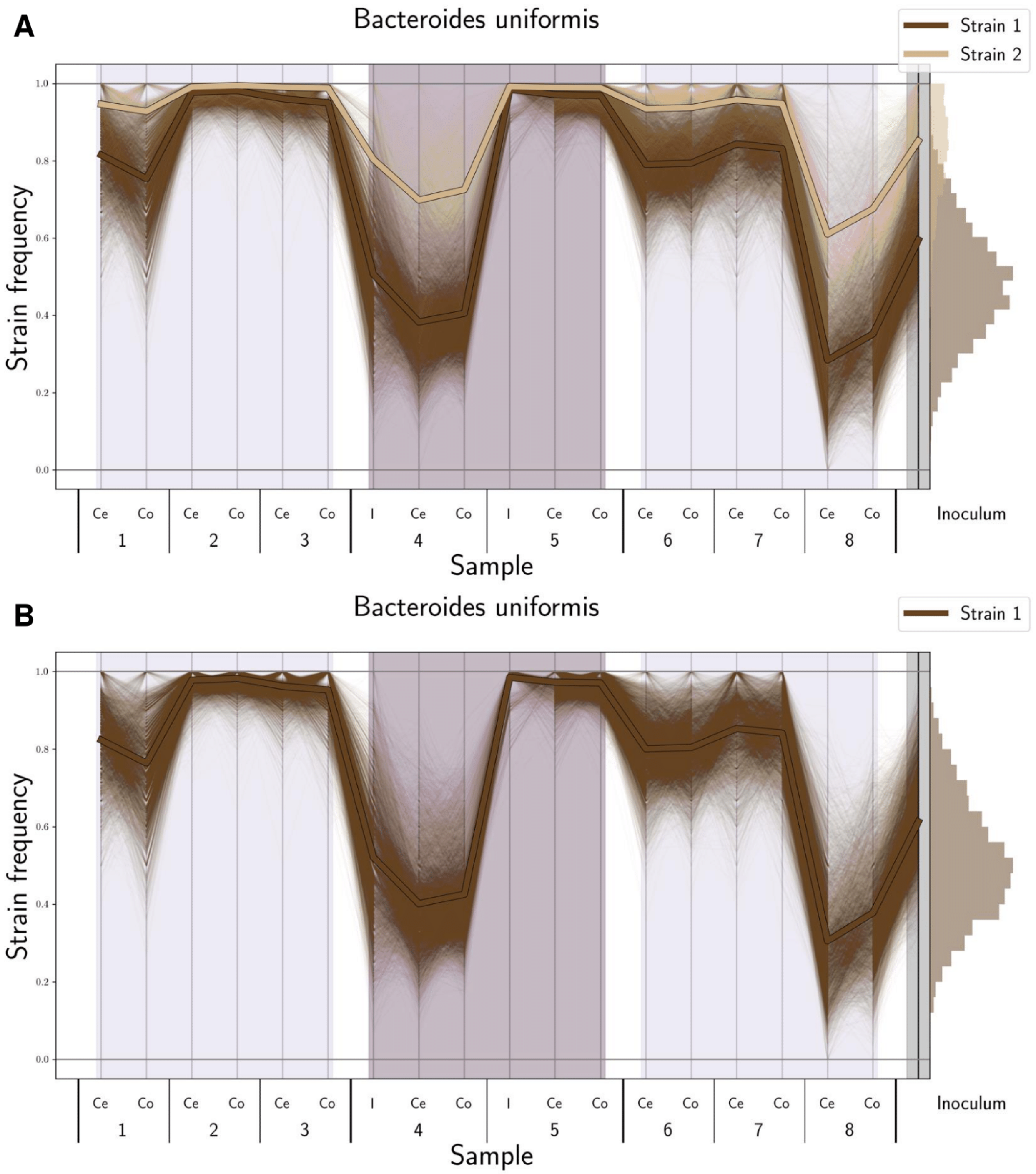


**Supplementary Figure 15. Supervised strain frequency inference of *Bacteroides uniformis* strains. (A)** *B. uniformis* SNVs were clustered into two groups, despite representing a single group of correlated SNVs. **(B)** The two clusters were merged, and SNVs that had a distance $d \leq 3.5$ with 25% of other SNVs in the cluster were retained.
